# Circuit mechanism underlying fragmented sleep and memory deficits in 16p11.2 deletion mouse model of autism

**DOI:** 10.1101/2023.12.26.573156

**Authors:** Ashley Choi, Jennifer Smith, Yingqi Wang, Hyunsoo Shin, Bowon Kim, Alyssa Wiest, Xi Jin, Isabella An, Jiso Hong, Hanna Antila, Steven Thomas, Janardhan P. Bhattarai, Kevin Beier, Minghong Ma, Franz Weber, Shinjae Chung

**Affiliations:** Department of Neuroscience, Chronobiology and Sleep Institute, Perelman School of Medicine, University of Pennsylvania, Philadelphia, PA 19104, USA; Department of Pharmacology, Perelman School of Medicine, University of Pennsylvania, Philadelphia, PA 19104, USA; Department of Physiology and Biophysics, School of Medicine, University of California, Irvine, Irvine, CA 92617, USA

## Abstract

Sleep disturbances are prevalent in children with autism spectrum disorder (ASD) and have a major impact on the quality of life. Strikingly, sleep problems are positively correlated with the severity of ASD symptoms, such as memory impairment. However, the neural mechanisms underlying sleep disturbances and cognitive deficits in ASD are largely unexplored. Here, we show that non-rapid eye movement sleep (NREMs) is highly fragmented in the 16p11.2 deletion mouse model of ASD. The degree of sleep fragmentation is reflected in an increased number of calcium transients in the activity of locus coeruleus noradrenergic (LC-NE) neurons during NREMs. Exposure to a novel environment further exacerbates sleep disturbances in 16p11.2 deletion mice by fragmenting NREMs and decreasing rapid eye movement sleep (REMs). In contrast, optogenetic inhibition of LC-NE neurons and pharmacological blockade of noradrenergic transmission using clonidine reverse sleep fragmentation. Furthermore, inhibiting LC-NE neurons restores memory. Rabies-mediated unbiased screening of presynaptic neurons reveals altered connectivity of LC-NE neurons with sleep- and memory regulatory brain regions in 16p11.2 deletion mice. Our findings demonstrate that heightened activity of LC-NE neurons and altered brain-wide connectivity underlies sleep fragmentation in 16p11.2 deletion mice and identify a crucial role of the LC-NE system in regulating sleep stability and memory in ASD.

## INTRODUCTION

Many children with ASD suffer from sleep disturbances including delayed sleep onset, frequent night awakenings, and short sleep episodes^1–3^. Strikingly, sleep disturbances worsen ASD symptoms, especially cognitive capabilities^4–6^. Yet, the neural mechanisms underlying sleep problems and the resulting cognitive impairment in ASD remain poorly understood.

Chromosomal copy number variations (CNVs) are associated with an increased prevalence of ASD; in particular, CNVs in chromosomal region 16p11.2 increase the risk for ASD^7–11^. 16p11.2 CNVs are associated with both sleep disturbances and cognitive impairment^11–13^. In particular, 16p11.2 hemideletion (16p11.2 del/+) mice display sleep disturbances and cognitive deficits^14–18^; the underlying mechanisms, however, are largely unknown.

Recent studies demonstrated that LC-NE neurons are rhythmically activated during NREMs in synchrony with an infraslow rhythm fluctuating on a ∼1-minute time-scale (infraslow σ rhythm), which is reflected in the electroencephalogram (EEG) σ power and the frequency of sleep spindles^19–21^. The activation of LC-NE neurons during NREMs tends to overlap with microarousals (MAs) while their activity decreases before transitions to REMs, suggesting that the infraslow oscillations of the LC-NE activity during NREMs coordinate the onset of MAs and REMs^19–24^. The activity of LC neurons and NE levels are increased during NREMs after learning, suggesting that the activation of LC-NE neurons during NREMs is involved in memory consolidation^25,26^. Activating LC-NE activity during post-learning sleep impairs spatial learning and memory performance, while suppressing their activity improves memory^21,23^. However, the extent to which LC-NE neurons contribute to sleep disturbances and memory deficits observed in ASD remains unknown.

By employing fiber photometry, optogenetic and pharmacological manipulations, spatial object recognition task, and mono-synaptically restricted rabies virus tracing, we investigated the role of LC-NE neurons in sleep disturbances and memory impairment in the 16p11.2 deletion mouse model of ASD.

## RESULTS

### 16p11.2 del/+ mice exhibit fragmented sleep and impaired phase-coupling of sound-evoked arousals with infraslow σ rhythm

We performed electroencephalogram (EEG) and electromyography (EMG) recordings in 16p11.2 del/+ mice and WT littermates to examine their sleep architecture throughout 24 hour recordings (**Fig. 1A**). The overall time spent in NREMs, REMs, and wakefulness was not significantly different between 16p11.2 del/+ mice and WT littermates (**Figs. 1B, S1A and S1B**; t tests, P = 0.995, 0.334, 0.388 for NREMs, REMs and wake; detailed statistical results are shown in **Table S1**). However, the number of MAs during NREMs was significantly higher in 16p11.2 del/+ mice, resulting in a decreased duration and increased frequency of NREMs episodes compared with WT mice (**Figs. 1B and 1C**; t tests, P = 0.020, 0.011, 0.009 for MAs, NREMs duration and frequency). MAs are defined as short (< 20 sec) wake-like episodes during sleep, characterized by desynchronized EEG and activated EMG^20,21,24,27–29^. Given their short duration, MAs have not been scored in previous sleep studies on 16p11.2 del/+ mice, which used a more coarse binning for sleep annotation than the 2.5 sec in our study^16,17^. Accordingly, examining the distribution of the duration of NREMs bouts, we found that 16p11.2 del/+ mice have an increased proportion of short NREMs bouts compared with WT mice (**Fig. 1D**; mixed ANOVA, duration P = 5.702e-40, genotype P = 1.000, interaction P = 7.109e-7). MAs preferentially occur at the descending phase of the infraslow σ rhythm, where mice are most susceptible to wake up in response to external stimulation^20,21,30,31^. We therefore examined the infraslow σ rhythm of 16p11.2 del/+ and WT mice. The strength of the infraslow σ rhythm was significantly decreased compared with that of WT mice (**Fig. 1E**; t test, P = 0.029). The number of sleep spindles, a major contributor to EEG σ power, was not affected in 16p11.2 del/+ mice (**Fig. S1C**; t test, P = 0.887), suggesting that decreased strength of infraslow σ rhythm in 16p11.2 del/+ is caused by a weakened clustering of sleep spindles. Finally, we confirmed that the sleep fragmentation and weakened infraslow σ rhythm phenotype in 16p11.2 del/+ mice was also conserved when they are crossed with dopamine β-hydroxylase (DBH)-Cre or GAD2-Cre mice used in the remainder of this study (**Figs. S1D-K**). Thus, 16p11.2 del/+ mice have an increased number of MAs resulting in sleep fragmentation, and their infraslow σ rhythm is weakened.

**Figure 1.**
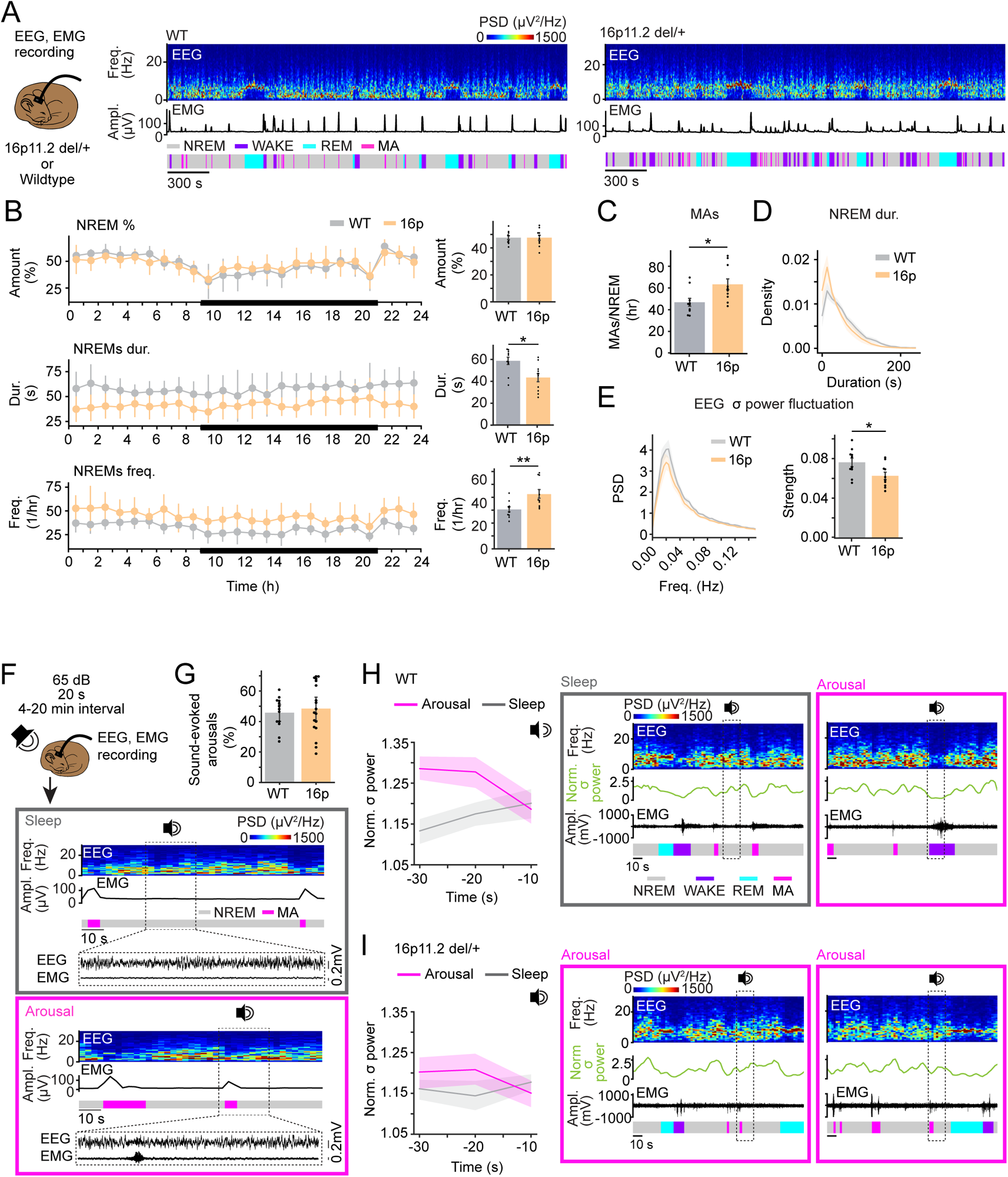
16p11.2 del/+ mice exhibit fragmented sleep and impaired phase-coupling of sound-evoked arousals with infraslow σ rhythm. **A.** Schematic of EEG and EMG recordings in 16p11.2 del/+ mice and WT littermates (left). Example sessions from WT and 16p11.2 del/+ mouse (middle and right). Shown are EEG power spectra, EMG amplitude, and color-coded sleep-wake states. **B.** Percentage of NREMs, duration and frequency of NREMs episodes during 24 hr recordings. Black line represents the dark phase. **C.** Number of MAs during NREMs. **D.** Distribution of NREMs episodes depending on their episode duration. **E.** Power spectral density (PSD) of normalized EEG σ power during NREMs. **F.** Schematic illustrating sound-evoked arousals. During about half of the trials, mice stayed asleep (gray box) or transitioned to wake (pink box) after sound. Shown are EEG power spectra, EMG amplitude, color-coded sleep-wake states, and EEG and EMG raw traces during selected periods (dotted box, sound stimuli). **G.** Percentage of sound-evoked arousals in WT and 16p11.2 del/+ mice. **H.** Left, EEG σ power before sound-evoked arousal and sleep-through trials in WT mice. Middle and right, example sleep-through trial with rising EEG σ power and wake-up trial with falling EEG σ power in a WT mouse. Shown are EEG power spectra, EEG σ power, EMG raw traces and color-coded sleep-wake states. Dotted box, sound stimuli. **I.** Left, EEG σ power before sound-evoked arousal and sleep-through trials in 16p11.2 del/+ mice. Middle and right, example wake-up trials with rising and falling EEG σ power in a 16p11.2 del/+ mouse. **B-E.** n = 10 WT and 10 16p11.2 del/+ mice. **F-I.** 16p11.2 del/+ mice crossed with B6129SF1/J mice or GAD2-Cre mice were used. n = 18 16p11.2 del/+ and 16p11.2 del/+ x GAD2-Cre mice and 13 WT and WTx GAD2-Cre mice. Bars and lines, averages across mice; dots, individual mice; error bars, s.e.m. T tests, **P < 0.01; *P < 0.05.

During NREMs, mice are more susceptible to wake up in response to acoustic stimuli during the descending phase of the infraslow σ rhythm, whereas they tend to sleep through during the ascending phase^30^. Because 16p11.2 del/+ mice exhibit a weakened infraslow σ rhythm, we investigated whether the phase tuning of sound-evoked arousals is altered in 16p11.2 del/+ and WT mice. Sounds (65 dB, 20 s) were randomly presented every 4-20 min, while performing sleep recordings in WT and 16p11.2 del/+ mice (**Fig. 1F**). Both groups of mice woke up during ∼ half of the trials in response to the tone, suggesting that the probability to be awakened by an external sound is not altered in 16p11.2 del/+ mice (**Fig. 1G**). In WT mice, the EEG σ power started slowly decreasing ∼ 20 s before the tone onset in sound-evoked arousal trials, while it increased in sleep-through trials (**Fig. 1H**; two-way repeated measures [rm] ANOVA, time P = 0.507, wake P = 0.040, interaction P = 0.033). In contrast, in 16p11.2 del/+ mice, the time course of the EEG σ power was similar in both arousal and sleep-through trials (**Fig. 1I**; two-way rm ANOVA, time P = 0.883, wake P = 0.370, interaction P = 0.248). Thus, 16p11.2 del/+ mice exhibit an impaired phase tuning of sound-evoked arousals with the infraslow σ rhythm.

### Increased activation of LC-NE neurons during NREMs in 16p11.2 del/+ mice

Previous studies showed that MAs during NREMs closely overlap with calcium transients in LC-NE neurons^20,21,24^. To test whether an increased number of calcium transients may contribute to the heightened MA frequency in 16p11.2 del/+ mice, we monitored the population activity of LC-NE neurons using fiber photometry. To express genetically encoded calcium indicators in LC-NE neurons, we crossed 16p11.2 del/+ mice with DBH-Cre mice and injected adeno-associated viruses (AAVs) with Cre-dependent expression of GCaMP6s (AAV-FLEX-GCaMP6s) into the LC of 16p11.2 del/+ x DBH-Cre or 16p11.2 +/+ (WT) x DBH-Cre mice (**Fig. 2A**). In both groups, the LC-NE neurons were highly active during wakefulness, less active during NREMs and almost silent during REMs (**Figs. 2B and 2C**; one-way rm ANOVA, P = 8.259e-29, 1.736e-21; pairwise t tests with Bonferroni correction, P = 7.631e-17, 9.336e-13 for REMs vs. Wake, 6.854e-15, 1.723e-11 for Wake vs. NREMs, 1.125e-11, 2.314e-8 for REMs vs. NREMs in WT and 16p). During NREMs, the calcium transients of LC-NE neurons were often accompanied by MAs (**Fig. 2B**). The number of LC-NE calcium transients and the probability that they coincide with MAs was significantly higher in 16p11.2 del/+ x DBH-Cre mice compared with WT x DBH-Cre mice (**Fig. 2D**; t tests, P = 0.001, 0.003 for Ca^2+^ transients and overlap with MAs), suggesting that the heightened LC-NE activity underlies the increased frequency of MAs and the resulting sleep fragmentation in 16p11.2 del/+ x DBH-Cre mice.

**Figure 2.**
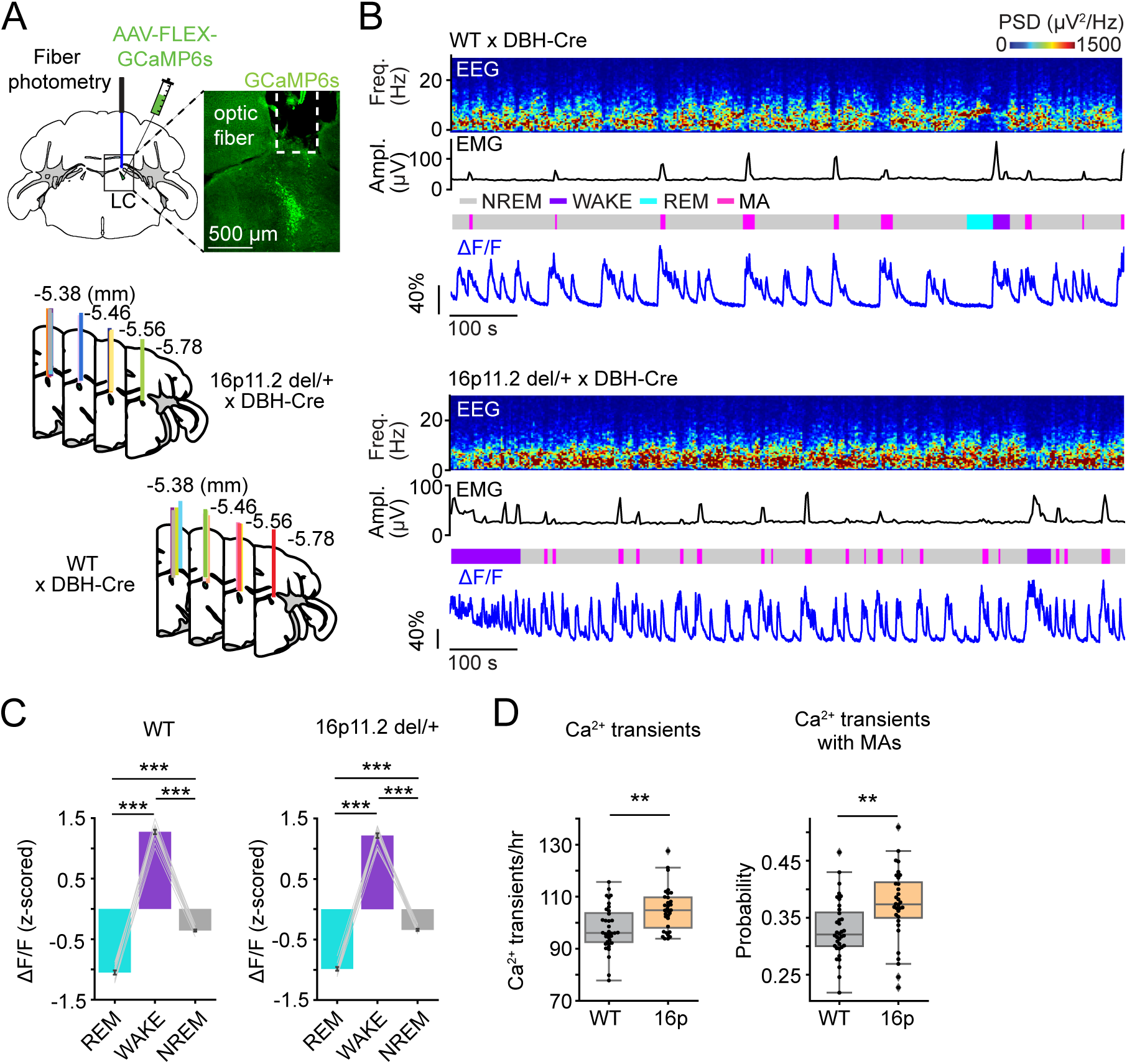
Increased activation of LC-NE neurons during NREMs in 16p11.2 del/+ mice. **A.** Schematic of fiber photometry with simultaneous EEG and EMG recordings in LC-NE neurons. Mouse brain figure adapted from the Allen Reference Atlas - Mouse Brain (atlas.brain-map.org). Top right, fluorescence image in a 16p11.2 del/+ x DBH-Cre mouse injected with AAV-FLEX-GCaMP6s into the LC. Scale bar, 500 μm. Bottom, location of optic fiber tracts. Each colored bar represents the location of optic fibers for photometry recordings. **B.** Example fiber photometry recordings of LC-NE neurons in a WT x DBH-Cre (top) and a 16p11.2 del/+ x DBH-Cre mouse (bottom). Shown are EEG power spectra, EMG amplitude, color-coded sleep-wake states, and ΔF/F signal. **C.** z-scored ΔF/F activity in WT x DBH-Cre and 16p11.2 del/+ x DBH-Cre mice. Bars, averages across mice; lines, individual mice; error bars, s.e.m. One-way rm ANOVA followed by pairwise t tests with Bonferroni correction, ***P < 0.001. **D.** Left, number of calcium transients in LC-NE neurons during NREMs. Right, proportion of calcium transients coinciding with MAs. Box plots; dots, individual mice. T tests, **P < 0.01. n = 12 WT x DBH-Cre and 11 16p11.2 del/+ x DBH-Cre mice.

### Elevated activity of LC-NE neurons in novel environment leads to fragmented NREMs and reduced REMs in 16p11.2 del/+ mice

In children with ASD, exposure to a novel environment worsens sleep efficiency^32,33^. As a behavioral paradigm to test the impact of a novel environment on sleep, we placed the same 16p11.2 del/+ and WT mice in a novel or familiar cage, while measuring their sleep using EEG/EMG recordings. To measure the overall sleep architecture and the activity of LC-NE neurons in a new or familiar environment, 16p11.2 del/+ x DBH-Cre mice and WT x DBH-Cre mice were injected with AAV-FLEX-GCaMP6s followed by an implantation of an optic fiber targeting the LC as well as EEG and EMG electrodes (**Fig. 3A**). The number of MAs during NREMs was significantly increased in 16p11.2 del/+ x DBH-Cre in the novel cage compared with the familiar cage resulting in decreased duration of NREMs episodes (**Figs. 3B and S2A**; mixed ANOVA, genotype P = 3.865e-4, 2.055e-5, condition P = 0.002, 0.002, interaction P = 0.005, 0.063; pairwise t tests with Holm correction, P = 0.003, 0.003 for MAs and NREMs duration). In contrast, WT x DBH-Cre mice did not change the level of MAs regardless of whether they were in the novel or familiar cage (**Fig. 3B**; P = 0.629). The level of MAs in the novel cage was significantly higher in 16p11.2 del/+ x DBH-Cre mice compared with WT x DBH-Cre mice (**Fig. 3B**; P = 0.001). In both groups, the amount of REMs was significantly decreased in the novel cage compared with the familiar cage, while the level was significantly lower in 16p11.2 del/+ x DBH-Cre mice compared with WTx DBH-Cre mice (**Fig. 3C**; mixed ANOVA, genotype P = 0.031, condition P = 1.333e-13, interaction P = 0.198; pairwise t tests with Holm correction, P = 0.029). In both groups of mice, exposure to the novel cage significantly increased the amount of wake while decreasing NREMs compared with the familiar cage (**Fig. 3C**; mixed ANOVA, condition P = 1.893e-7, 1.998e-5 for wake and NREMs). Moreover, we found that the number of LC-NE calcium transients and their overlap with MAs was significantly higher in 16p11.2 del/+ x DBH-Cre mice than in WT x DBH-Cre mice in the novel cage (**Figs. 3D and 3E**; t tests P = 0.030 and 0.014). The number of calcium peaks was positively correlated with the number of MAs and negatively correlated with the amount of REMs (**Fig. 3F**; linear regression r^2^ = 0.624, 0.334, P = 3.418e-5, 0.008 for correlation with MAs and REMs). Thus, elevated activity of LC-NE neurons in 16p11.2 del/+ x DBH-Cre mice following exposure to a novel cage leads to increased MAs during NREMs and reduced REMs.

**Figure 3.**
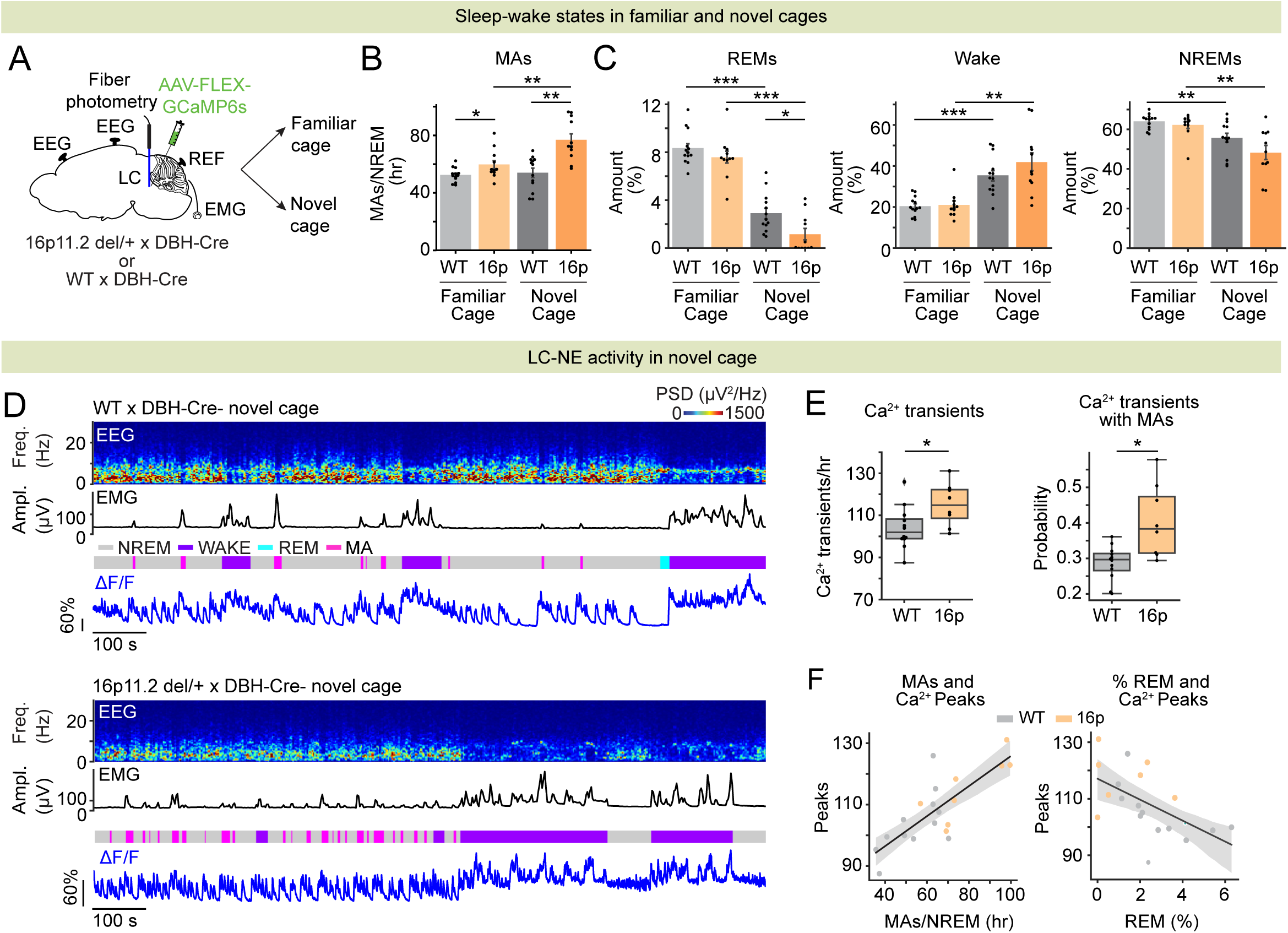
Elevated activity of LC-NE neurons in novel environment leads to fragmented NREMs and reduced REMs in 16p11.2 del/+ mice. **A.** Schematic of fiber photometry with simultaneous EEG and EMG recordings in LC-NE neurons during exposure to a familiar or novel cage. **B.** Number of MAs during NREMs in 16p11.2 del/+ x DBH-Cre and WT x DBH-Cre mice in the novel and familiar cages. **C.** Percentage of time spent in REMs, wakefulness and NREMs. **D.** Example fiber photometry recordings of LC-NE neurons in a WT x DBH-Cre (top) and a 16p11.2 del/+ x DBH-Cre mouse (bottom) in a novel cage. Shown are EEG power spectra, EMG amplitude, color-coded sleep-wake states, and ΔF/F signal. **E.** Left, number of calcium transients in LC-NE neurons during NREMs. Right, proportion of calcium transients coinciding with MAs. Box plots; dots, individual mice. T tests, *P < 0.05. **F.** Correlation of the number of calcium transients with the number of MAs (left) or the amount of REMs (right). Shadings, 95% CIs. **B-C.** Mixed ANOVA followed by pairwise t tests with Holm correction. ***P < 0.001; **P < 0.01; *P < 0.05 n = 11 16p11.2 del/+ x DBH-Cre and 13 WT x DBH-Cre mice.

### Optogenetic inhibition of LC-NE neurons and clonidine reverses sleep fragmentation in 16p11.2 del/+ mice

To test whether inhibiting LC-NE neurons reverses sleep fragmentation in 16p11.2 del/+ mice, we bilaterally injected AAVs encoding the bistable chloride channel SwiChR++ (AAV-DIO-SwiChR++-eYFP) or eYFP (AAV-DIO-eYFP) into the LC of 16p11.2 del/+ x DBH-Cre mice to optogenetically inhibit LC-NE neurons (**Figs. 4A and 4B**). We performed sustained optogenetic inhibition (3 s step pulses at 3 min intervals for 6 hrs) and compared the number of MAs as well as the percentage, duration, and frequency of each state. SwiChR++-mediated inhibition of LC-NE neurons significantly decreased the number of MAs, while increasing and decreasing the duration and frequency of NREMs episodes, respectively (**Figs. 4C and 4D**; t tests, P = 2.938e-4, 0.377, 0.002, 6.865e-4 for MAs, NREM %, duration and frequency). The overall time spent in NREMs, REMs, and wakefulness was not changed in both groups (**Figs. 4D, S3A and S3B**; t tests, P > 0.161). Optogenetic inhibition significantly decreased the peak frequency of the infraslow rhythm (**Fig. S3C**; 0.015 ± 0.005 Hz in eYFP and 0.006 ± 0.002 Hz in SwiChR; t test, P = 5.198e-5), which may contribute to reduced number of MAs. Thus, inhibiting LC-NE neurons in 16p11.2 del/+ x DBH-Cre mice reversed sleep fragmentation without changing the overall amount of sleep.

**Figure 4.**
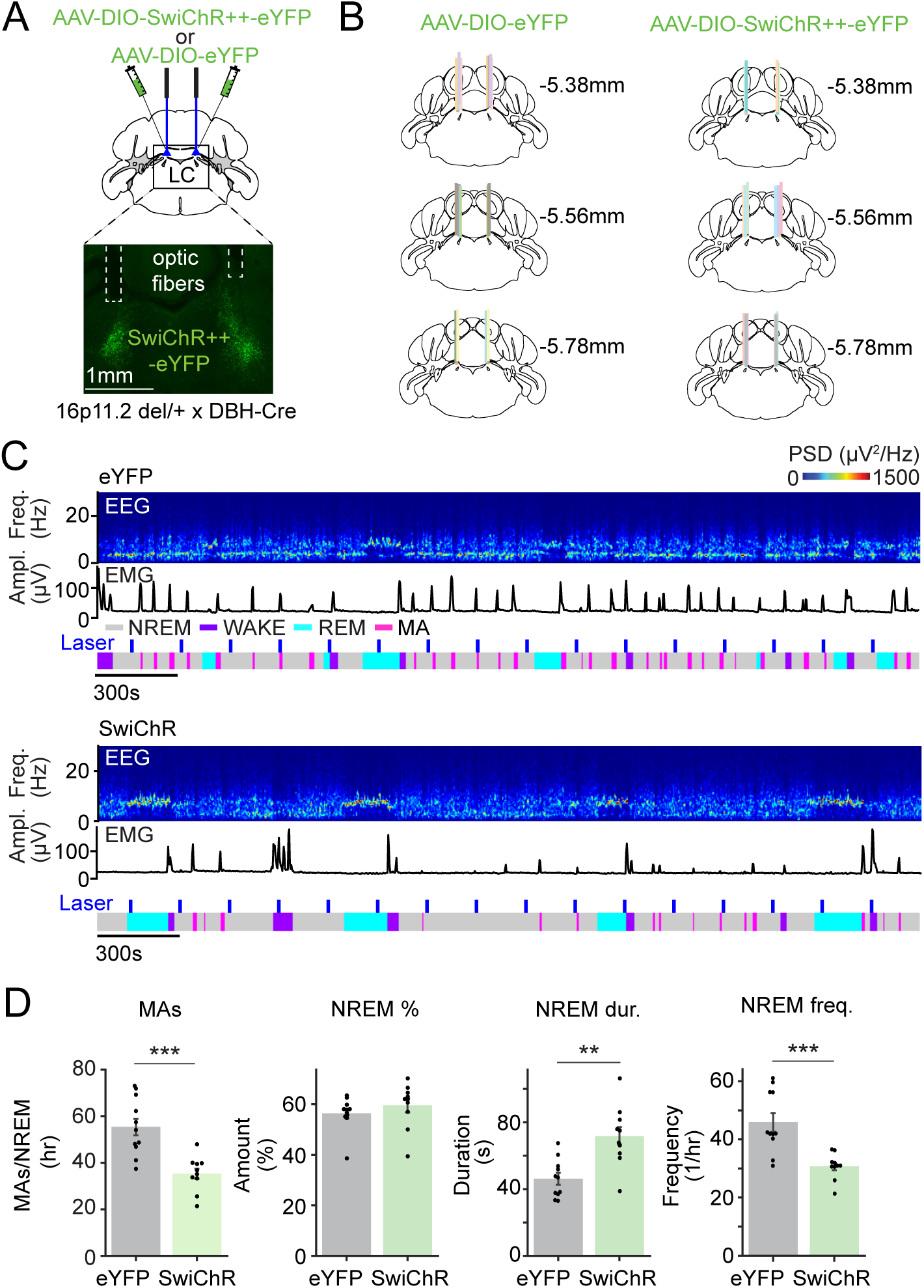
Optogenetic inhibition of LC-NE neurons reverses sleep fragmentation in 16p11.2 del/+ mice. **A.** Schematic of SwiChR++-mediated inhibition experiments and a fluorescence image of LC in a 16p11.2 del/+ x DBH-Cre mouse bilaterally injected with AAV-DIO-SwiChR++-eYFP into the LC. **B.** Location of fiber tracts. Each colored bar represents the location of optic fibers. **C.** Example sessions from eYFP- (top) or SwiChR++- (bottom) expressing 16p11.2 del/+ x DBH-Cre mouse with laser stimulation. Shown are EEG power spectra, EMG traces, and color-coded sleep-wake states. **D.** Number of MAs during NREMs, percentage of NREMs, duration and frequency of NREMs episodes in eYFP- or SwiChR++-expressing 16p11.2 del/+ x DBH-Cre mice during the 6 hr laser recordings. n = 11 eYFP-16p11.2 del/+ x DBH-Cre and 10 SwiChR-16p11.2 del/+ x DBH-Cre mice.

To test whether pharmacologically blocking noradrenergic transmission in 16p11.2 del/+ mice also reverses sleep fragmentation, we next tested the effect of clonidine, an ɑ2 adrenergic receptor (ADRA2A) agonist in 16p11.2 del/+ mice (1 mg/kg, intraperitoneal). Injection of clonidine significantly decreased the number of MAs while increasing the duration of NREMs episodes (**Figs. S4A-C**; paired t tests, P = 0.002, 0.019 for MAs and NREMs duration). The overall time spent in NREMs was also significantly increased, while wakefulness and REMs were decreased (**Figs. S4C-E**; P = 0.001, 0.027, 1.230e-5 for NREMs, wake, and REMs %). These effects on the amount of NREMs, REMs and wakefulness were different from those observed following optogenetic LC-NE inhibition. This may be caused by the fact that the pharmacological blockade of presynaptic ADRA2A using clonidine decreases overall noradrenaline release from both central and peripheral nerve terminals whereas optogenetic inhibition specifically inhibits the activity of noradrenergic cell bodies within the LC. Taken together, both optogenetic inhibition of LC-NE neurons and pharmacological blockade of noradrenergic transmission improved sleep continuity, while clonidine further increased the amount of NREMs and decreased wakefulness and REMs.

### Inhibiting LC-NE neurons restores memory in 16p11.2 del/+ mice

Previous studies showed that 16p11.2 del/+ mice exhibit impaired memory^14,15,18,34^. We tested whether 16p11.2 del/+ mice exhibit impaired memory in spatial object recognition (SOR) task compared with WT mice (**Fig. 5A**). Mice were first habituated to the box on the first two days. During the training session on the third day, mice were placed in the arena with two identical objects, and then returned to their home cage for 5 hrs. During the test session, mice were placed in the arena with the two familiar objects, one displaced to a new location. WT mice exhibited a significant increase in the preference for the moved object, whereas in 16p11.2 del/+ mice the memory was impaired (**Fig. 5B**; paired t tests, P = 0.025, 0.250 in WT and 16p mice; t test, P = 0.020 for discrimination ratio). To test whether inhibiting the activity of LC-NE neurons after the training session reverses memory deficits in 16p11.2 del/+ mice, we performed SwiChR++-mediated optogenetic inhibition of LC-NE neurons in 16p11.2 del/+ x DBH-Cre and WT x DBH-Cre mice (**Fig. 5C**). Immediately following the training session, optogenetic inhibition was performed for 5 hours in the home cage as described above (3 s step pulses at 3 min intervals). WT mice still showed a significant preference for the moved object after inhibition of LC-NE neurons, and the change in the preference between the training and test session was comparable for eYFP - WT x DBH-Cre and SwiChR - WT x DBH-Cre groups (**Fig. 5D**; paired t tests, P = 0.018, 0.049 in eYFP and SwiChR mice; t test, P = 0.254 for discrimination ratio). However, in 16p11.2 del/+ x DBH-Cre mice, SwiChR++-mediated inhibition of LC-NE neurons significantly increased the preference for the moved object compared with the eYFP-expressing 16p11.2 del/+ x DBH-Cre control mice (**Fig. 5E**; paired t tests, P = 0.382, 0.002 in eYFP and SwiChR mice; t test, P = 0.017 for discrimination ratio). Optogenetic inhibition after the training in 16p11.2 del/+ x DBH-Cre mice also decreased the amount of MAs, similar to Fig. 4D (**Fig. S5A**; t test, P = 0.048). Taken together, inhibiting LC-NE neurons reversed memory impairment in 16p11.2 del/+ mice.

**Figure 5.**
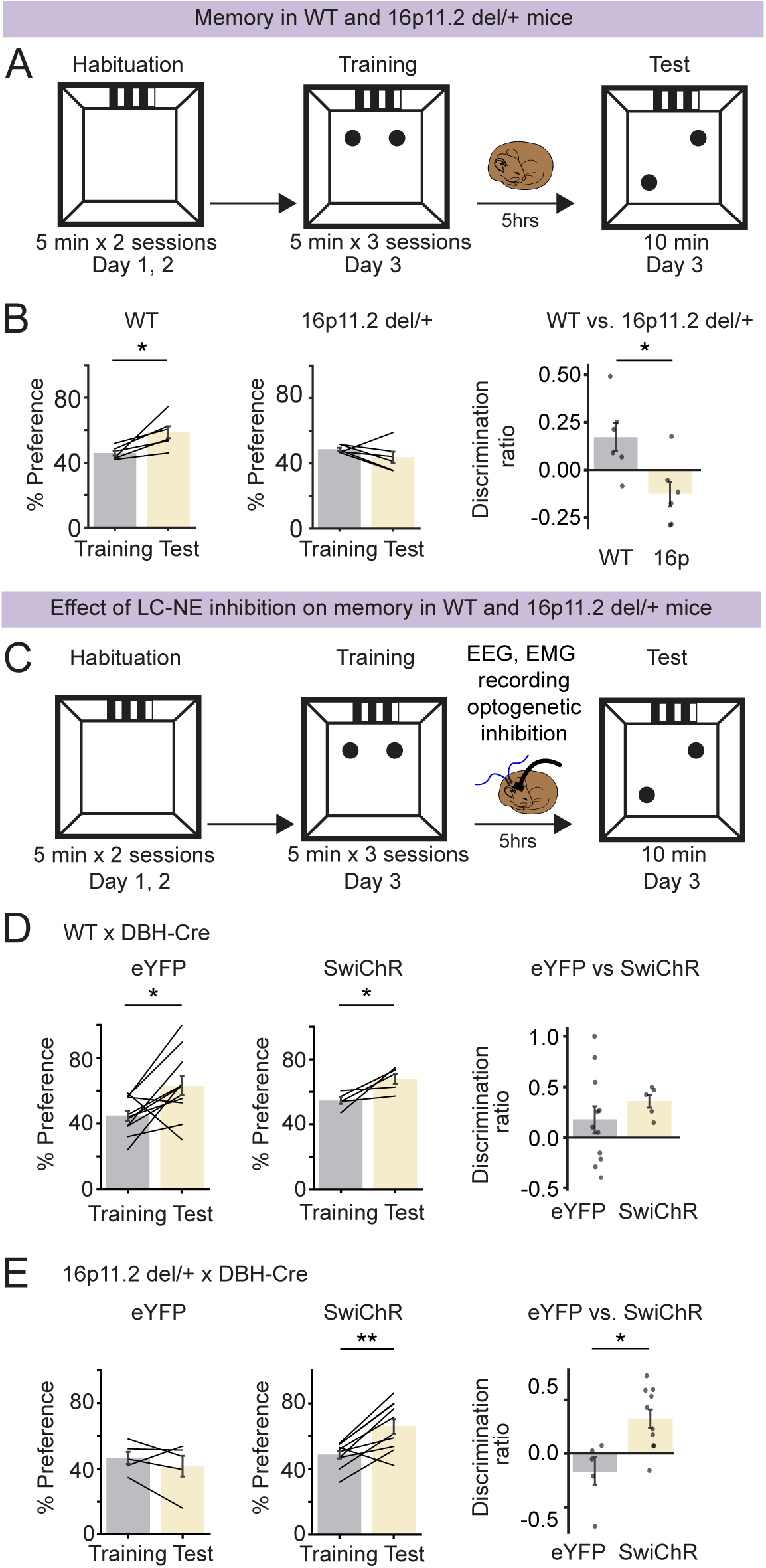
Optogenetic inhibition of LC-NE neurons restores memory in 16p11.2 del/+ mice. **A.** Schematic of SOR task in WT and 16p11.2 del/+ mice. **B.** Preference (%) for a moved object during training and testing sessions in WT and 16p11.2 del/+ mice (left and middle). Preference (%) is calculated as 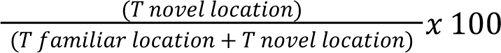. Discrimination ratio (right) is calculated as 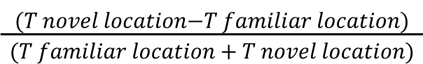. n = 6 WT and 6 16p11.2 del/+ mice. Schematic of SOR task combined with SwiChR++-mediated LC-NE inhibition. **C.** Effect of LC-NE inhibition on SOR memory in WT x DBH-Cre mice. n = 11 eYFP-WT x DBH-Cre and 5 SwiChR - WT x DBH-Cre mice. **D.** Effect of LC-NE inhibition on SOR memory in 16p11.2 del/+ x DBH-Cre mice. n = 5 eYFP-16p11.2 del/+ x DBH-Cre and 10 SwiChR-16p11.2 del/+ x DBH-Cre mice. Bars, averages across mice; dots and lines, individual mice; error bars, s.e.m. Unpaired and paired t tests, **P < 0.01; *P < 0.05.

### Changes of LC-NE presynaptic input distributions in 16p11.2 del/+ mice

Differences in the distribution of presynaptic inputs may contribute to the different activity dynamics of LC-NE neurons in 16p11.2 del/+ and WT mice. To compare the organization of presynaptic inputs between 16p11.2 del/+ and WT mice, we performed mono-synaptically restricted rabies virus tracing from LC-NE neurons. AAVs with Cre-dependent expression of the TVA receptor fused with mCherry (TC66T) and rabies glycoprotein (RG) were injected into the LC of 16p11.2 del/+ x DBH-Cre and WT x DBH-Cre mice (**Fig. 6A**). 12 days later, a modified rabies virus expressing GFP (RV*d*G-GFP) was injected into the LC. The majority of starter cells expressing both TC66T and GFP were located in the LC (**Fig. 6A**). The proportion of input neurons in the hypothalamus was similar in both groups (**Fig. 6B**; bootstrap, P = 0.48). However, the proportion in the lateral hypothalamic area (LHA) was higher in 16p11.2 del/+ mice compared with WT mice (**Fig. 6C**; P = 0.012). We have recently shown that LHA glutamatergic neurons are wake- and MA-promoting^35^. While the cell type of these presynaptic inputs in the LHA remains to be determined, we predict that the increased number of inputs from the LHA to the LC in 16p11.2 del/+ mice may contribute to increased frequency of MAs. The proportion of input neurons in the medulla of 16p11.2 del/+ mice was higher compared with WT mice (**Fig. 6B**; P = 0.042). In particular, we found more presynaptic cells in the ventral medulla (VM) of 16p11.2 del/+ mice (**Fig. 6C**; P < 0.001). The VM sends excitatory inputs to the LC^36^. Activation of glutamatergic neurons in the VM strongly promotes wakefulness^37^, and these inputs to the LC may lead to increased arousal in 16p11.2 del/+ mice. A small proportion of input neurons was found in the cortex and hippocampus of WT mice, and their number was further reduced in 16p11.2 del/+ mice (**Figs. 6B and 6C**; P < 0.001).

**Figure 6.**
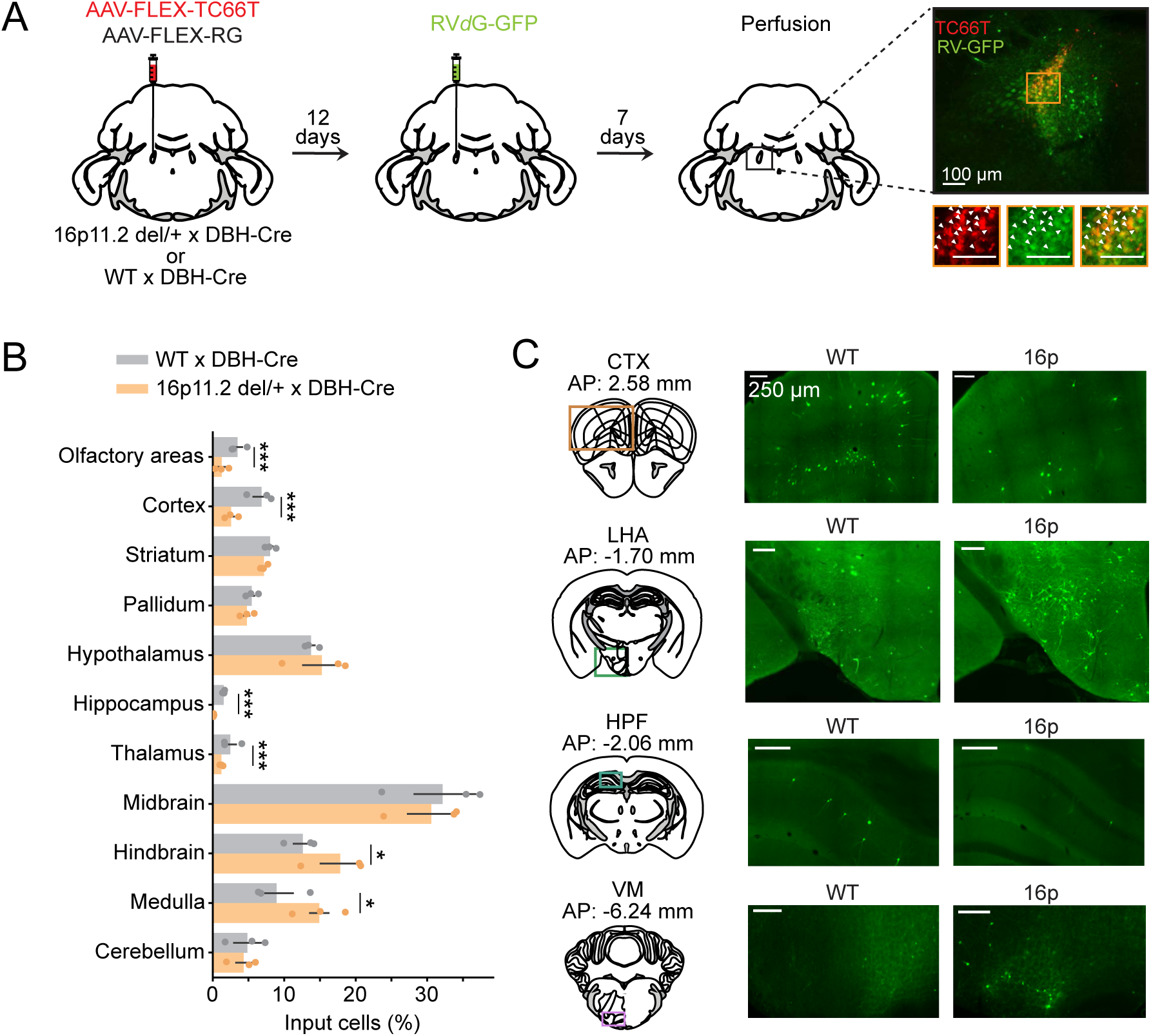
Changes of LC-NE presynaptic input distributions in 16p11.2 del/+ mice. **A.** Schematic illustration of rabies-mediated tracing of monosynaptic inputs to LC-NE neurons. AAVs expressing Cre-dependent mutant EnvA receptor fused with mCherry (TC66T) and Cre-dependent rabies glycoprotein (GP) were injected into the LC of 16p11.2 del/+ x DBH-Cre and WT x DBH-Cre mice. 12 days later, we injected EnvA-pseudotyped, G-deleted, and GFP expressing rabies virus (RV*d*G-GFP). Starter cells co-express TC66T and GFP (right). Scale bar, 100 μm. **B.** Proportion of RV-GFP labeled inputs to LC-NE neurons across brain regions defined by the Allen Reference Atlas - Mouse Brain (atlas.brain-map.org). Proportion of input cells (%) was calculated by dividing the number of RV-GFP labeled neurons found in a specific brain region by the total number of input neurons. Bars, average across mice; dots, individual mice; error bars, s.e.m. n = 3 mice. Bootstrap, ***P < 0.001; *P < 0.05 **C.** GFP-labeled input neurons in the cortex (CTX), lateral hypothalamic area (LHA), hippocampus (HPF) and ventral medulla (VM, the shown image contains the magnocellular reticular nucleus and paragigantocellular reticular nucleus within the medulla) of 16p11.2 del/+ x DBH-Cre and WT x DBH-Cre mice. Scale bar, 250 μm.

To examine whether the increased activity of LC-NE neurons is due to the changes in their intrinsic excitability, we performed whole-cell patch clamp recordings of LC-NE neurons in 16p11.2 del/+ x DBH-Cre and WT x DBH-Cre mice. We injected AAV-DIO-tdTomato into the LC of these mice. Three weeks later, we performed patch clamp recordings of tdTomato expressing LC-NE cells. We found that LC-NE neurons in 16p11.2 del/+ x DBH-Cre mice display slightly lower firing frequencies upon current injection than WT x DBH-Cre mice (**Fig. S6**; two-way ANOVA, P = 0.022 for cell type).

Consequently, changes in the presynaptic inputs and not an increase in their intrinsic excitability likely underlie the increased activation of LC-NE neurons during NREMs.

## DISCUSSION

We have identified a neural mechanism underlying sleep fragmentation and memory deficits in the 16p11.2 deletion mouse model of ASD. We found that 16p11.2 del/+ mice exhibit frequent MAs, a weakened infraslow σ rhythm and impaired infraslow phase coupling of sound-evoked arousals (**Fig. 1**). The frequent MAs in 16p11.2 del/+ mice are reflected in an increased number of LC-NE calcium transients during NREMs (**Fig. 2**). Exposure to a novel environment further disrupted their sleep quality (**Fig. 3**). In contrast, optogenetic inhibition of LC-NE neurons and pharmacological blockade of noradrenergic transmission using clonidine reversed sleep fragmentation (**Fig. 4**). In addition, inhibition of LC-NE neurons restored memory in 16p11.2 del/+ mice (**Fig. 5**). Rabies virus-mediated screening revealed an altered connectivity of LC-NE neurons with presynaptic partners in 16p11.2 del/+ mice that may contribute to the observed sleep disturbances (**Fig. 6**). In sum, our findings elucidate a mechanism by which heightened LC-NE neuron activity underlies sleep disturbances and memory impairment in the 16p11.2 del mouse model of ASD.

In addition to the NE system, previous studies also found disturbances in the serotonergic system, causing hyperactivity, reduced sociability and difficulties in coping with acute stress in 16p11.2 del/+ mice ^38,39^. Serotonergic neurons in the dorsal raphe nucleus are also activated with the infraslow σ rhythm during NREMs^40^. For future studies, it would be interesting to test whether the activity of serotonergic neurons in 16p11.2 del/+ mice is also disturbed during sleep, possibly contributing to sleep disturbances and whether restoring sleep can alleviate hyperactivity and improve sociability. Besides NE and serotonin, dysfunctional dopaminergic signaling has also been shown to cause sleep disturbances. A homozygous InsG3680 mutation in the *Shank3* gene caused a reduction of NREMs^41^. Opto- and chemogenetic inhibition of ventral tegmental area dopaminergic neurons during the critical developmental period increased NREMs and restored social novelty preference in adulthood. For future studies, it would be interesting to test whether these deficits in dopaminergic signaling are specific to Shank3 mutant mice or whether different ASD mouse models including 16p11.2 del/+ mice share dysfunctions in these neuromodulatory systems.

Consistent with previous reports^14,15,18,34^, we observed an impairment in SOR memory in 16p11.2 del/+ mice. Inhibiting LC-NE activity during subsequent sleep to reduce MAs and consolidate sleep, improved memory. A previous study found reduced activation of LC-NE neurons in 16p11.2 del/+ mice during motor learning, and activating LC-NE neurons during rotating disk tasks improved motor learning^42^. Thus, while increased LC-NE activity during motor learning is beneficial, our study, consistent with previous studies^21,23^, implies that reducing heightened LC activity during subsequent sleep improves learning, likely as a result of improved sleep quality.

Given that the rate of sound-evoked arousals in 16p11.2 del/+ mice is similar to that in WT mice (**Fig. 1G**), their sleep disturbances are likely the result of internal processes causing the heightened frequency of MAs, rather than caused by external stimuli. It is generally assumed that fragmented sleep in individuals with ASD is due to an increased sensitivity to the external stimuli such as sound^43^. While we observed highly fragmented NREMs in 16p11.2 del/+ mice, our results contradict this assumption. The likelihood that WT and 16p11.2 del/+ mice wake up to sound is similar, but the phase coupling of the awakenings with the infraslow σ rhythm is impaired. The infraslow σ rhythm is thought to partition sleep into a consolidated phase, ideal for sleep-dependent processes such as memory consolidation, followed by a fragile sleep phase, where the organism is sensitive to external stimuli and can be easily awakened ^30^. Disruption in the infraslow σ rhythm may consequently interfere with internal memory processes, resulting in poor memory consolidation. In humans, an enhanced infraslow σ rhythm, in particular within the fast spindle frequency band, was shown to be associated with better overnight memory consolidation^30^. Our rabies virus tracing experiments indicate that the number of presynaptic input neurons from the hippocampus to the LC is decreased in 16p11.2 del/+ mice which may contribute to their memory impairment. LC projections to the hippocampus are crucial for learning and memory^44–47^, and the reduced number of LC-projecting neurons in the hippocampus may impair a feedback mechanism, possibly contributing to the memory deficits in 16p11.2 del/+ mice.

In children with ASD, exposure to a novel environment worsens sleep efficiency^32,33^, and similarly the exposure to a novel cage exacerbated sleep quality in 16p11.2 del/+ mice, reflected in reduced REMs and a tendency towards less NREMs compared with WT mice (**Figs. 3B and 3C**). 16p11.2 del/+ mice took longer time to habituate to their sleep environment, and the degree of habituation may therefore significantly impact the amount and quality of sleep in mouse models of ASD^16,17^. Our findings further imply that blocking noradrenergic transmission can not only improve sleep in ASD patients^48^, but help them to more quickly habituate to new sleep environments.

Taken together, our results demonstrate that heightened activity of LC-NE neurons contribute to fragment sleep and impaired memory in the 16p11.2 del mouse model of ASD. Elucidating the circuit mechanisms underlying various aspects of sleep disturbances will provide valuable insights for the development of targeted therapeutic interventions to specifically improve distinct features of sleep and related behaviors in ASD.

## MATERIALS and METHODS

### Animals

16p11.2 del/+ mice^49^, B6129SF1/J and GAD2-Cre mice were obtained from Jackson lab (#013128, 101043, 010802; Jackson lab). DBH-Cre mice^20,50^ were obtained from Dr. Steve Thomas, Penn. 16p11.2 del/+ mice were crossed with B6129SF1/J mice to generate 16p11.2 del/+ and 16p11.2 +/+ (WT) mice. To target LC-NE neurons for photometry recordings, optogenetic inhibition, RV tracing and patch clamp recordings, 16p11.2 del/+ mice were crossed with DBH-Cre mice to generate 16p11.2 del/+ x DBH-Cre and 16p11.2 +/+ (WT) x DBH-Cre mice. For sound stimulation experiments, 16p11.2 del/+ crossed with GAD2-Cre mice as well as 16p11.2 del/+ mice crossed with B6129SF1/J mice were used. All mice were housed on a 12 h light/12 h dark cycle (lights on 07:00 and off 19:00) with free access to food and water and were 6-16 weeks old at the time of surgery. A previous study reported male-specific sleep deficits in the 16p11.2 del/+ mice^16^ and therefore male mice were used for all the experiments. All procedures were approved by Institutional Animal Care and Use Committees of the University of Pennsylvania and were done in accordance with the federal regulations and guidelines on animal experimentation (National Institutes of Health Offices of Laboratory Animal Welfare Policy).

### Surgery

To implant electroencephalogram (EEG) and electromyogram (EMG) recording electrodes, mice were anesthetized with 1.5-2% isoflurane and placed on a stereotaxic frame. Two stainless steel screws were inserted into the skull 1.5 mm from midline and 1.5 mm anterior to the bregma, and 2.5 mm from midline and 2.5 mm posterior to the bregma. The reference screw was inserted on top of the cerebellum. Two EMG electrodes were inserted into the neck musculature. Insulated leads from the EEG and EMG electrodes were soldered to a 2 × 3 pin header, which was secured to the skull using dental cement.

For LC photometry recordings, AAV_1_-Syn-Flex-GCaMP6S-WPRE-SV40 (0.3 μl, #100845-AAV1, Penn vector core/Addgene) was injected into LC (AP -5.3 mm, ML +/-1.0 mm, DV -2.7-3.3 mm from the cortical surface) using Nanoject II (Drummond Scientific) via a micropipette followed by an optic fiber (400 µm in diameter) implantation on top of the injection site. For optogenetic inhibition experiments, AAV2-EF1α-DIO-eYFP or AAV2-EF1α-DIO-SwiChR++-eYFP (0.3 μl, #27056, 55631, University of North Carolina vector core) was bilaterally injected into the LC followed by bilateral implantation of optic fibers (200 μm in diameter). Dental cement was applied to cover the exposed skull completely and to secure the optic fiber and the EEG/EMG implant. After surgery, mice were allowed to recover for at least 2-3 weeks before experiments.

For patch clamp recordings, AAV-DIO-tdTomato (0.3 ul, #28306-AAV2, Addgene) was injected into LC. 3 weeks later, patch clamp recordings were performed.

For rabies tracing experiments, 0.4 μl of a mixture containing AAV-CAG-FLEX-TC66T and AAV-CAG-FLEX-RG was injected into the LC. After 12 days, 0.4 μl of EnvA-pseudotyped, rabies-glycoprotein-deleted, and GFP-expressing rabies viral particles (RV*dG*) were injected into the LC. After 7 days, mice were perfused. Viruses for rabies tracing experiments were obtained from the University of California, Irvine.

### Histology

Mice were deeply anesthetized and transcardially perfused with phosphate-buffered saline (PBS) followed by 4% paraformaldehyde (PFA) in PBS. Brains were fixed overnight in 4% PFA and then transferred to 30% sucrose in PBS solution for at least one night. Brains were embedded and mounted with Tissue-Tek OCT compound (Sakura Finetek) and frozen. 50-60 μm sections were cut using a cryostat (Thermo Scientific HM525 NX) and mounted onto glass slides. Brain sections were washed in PBS, permeabilized using PBST (0.3% Triton X-100 in PBS) for 30 minutes and then incubated in blocking solution (5% normal donkey serum [NDS] in PBST) for 1 hour. Slices were then incubated in primary antibodies (1:1000, anti-GFP antibody, #GFP879484, Fisher Scientific) overnight at 4 °C. 18-24 hrs later, slices were washed with PBS and incubated with the corresponding secondary antibody (1:500; Alexa Fluor 488-AffiniPure donkey anti-chicken IgY (IgG) (H+L), #703545155, Jackson ImmunoResearch Laboratories). Brain sections were washed with PBS followed by counterstaining with Hoechst solution (#33342, Thermo Scientific) and mounted onto glass slides. Slides were cover-slipped with Fluoromount-G (Southern Biotechnic) and imaged using a fluorescence microscope (Microscope, Leica DM6B; Camera, Leica DFC7000GT; LED, Leica CTR6 LED).

### Sleep recordings

Sleep recordings were carried out in the animal’s home cage or in a cage to which the mouse had been habituated. EEG and EMG electrodes were connected to flexible recording cables via a mini-connector. EEG and EMG signals were recorded using an RHD2132 amplifier (Intan Technologies, sampling rate 1 kHz) connected to the RHD USB Interface Board (Intan Technologies). For fiber photometry, we used a Tucker-Davis Technologies RZ5P amplifier (sampling rate 1.5 kHz). EEG and EMG signals were referenced to a ground screw placed on top of the cerebellum. To determine the sleep-wake state of the animal, we first computed the EEG and EMG spectrogram for sliding, half-overlapping 5 s windows, resulting in 2.5 s time resolution. To estimate within each 5 s window the power spectral density (PSD), we performed Welch’s method with Hanning window using sliding, half-overlapping 2 s intervals. Next, we computed the time-dependent δ (0.5 to 4 Hz), θ (5 to 12 Hz), σ (12 to 20 Hz) and high γ (100 to 150 Hz) power by integrating the EEG power in the corresponding ranges within the EEG spectrogram. We also calculated the ratio of the θ and δ power (θ/δ) and the EMG power in the range 50 to 500 Hz. For each power band, we used its temporal mean to separate it into a low and high part (except for the EMG and θ/δ ratio, where we used the mean plus one standard deviation as threshold). REMs was defined by a high θ/δ ratio, low EMG, and low δ power. A state was set as NREMs if the δ power was high, the θ/δ ratio was low, and EMG power was low. In addition, states with low EMG power, low δ, but high σ power were scored as NREMs. Wake encompassed states with low δ power and high EMG power and each state with high γ power (if not otherwise classified as REMs). Our automatic algorithm has been published^20,35,37,51–53^ and has 90.256 % accuracy compared with the manual scoring by expert annotators. We manually verified the automatic classification using a graphical user interface visualizing the raw EEG and EMG signals, EEG spectrograms, EMG amplitudes, and the hypnogram to correct for errors, by visiting each single 2.5 sec epoch in the hypnograms. The software for automatic sleep-wake state classification and manual scoring was programmed in Python (available at https://github.com/tortugar/Lab/tree/master/PySleep).

### Sound-evoked arousal

Mice were exposed to acoustic stimuli (15000 Hz, 65 dB, 20 sec) randomly every 4-20 min for 6-7 hrs of recordings. The duration of noise and the sound level was based on the arousal success rate that mice woke up or slept through half of trials (arousal success rate was ∼ 50%). For analysis, episodes with NREMs duration > 120 sec were used for analysis. All sound trials were divided into sleep-through and arousal trials and EEG σ power was measured before the onset of sound.

### Optogenetic manipulation

Light pulses (3 s step pulses at 3 min intervals, 2 - 4 mW) were generated by a blue laser (473 nm, Laserglow) and sent through the optic fiber (200 µm diameter, ThorLabs) that connects to the ferrule on the mouse head. TTL pulses to trigger the laser were controlled using a raspberry pi, which was controlled by a custom user interface programmed in Python. Optogenetic manipulations were conducted during the light period for 6 hrs in Fig. 4 and 5 hrs in Fig. 5.

### Fiber photometry

For calcium imaging, a first LED (Doric lenses) generated the excitation wavelength of 465 nm and a second LED emitted 405 nm light, which served as control for bleaching and motion artifacts. The 465 and 405 nm signals were modulated at two different frequencies (210 and 330 Hz). Both lights were passed through dichroic mirrors before entering a patch cable attached to the optic fiber. Fluorescence signals emitted by GCaMP6s were collected by the optic fiber and passed via the patch cable through a dichroic mirror and GFP emission filter (Doric lenses) before entering a photoreceiver (Newport Co.). Photoreceiver signals were relayed to an RZ5P amplifier (Tucker-Davis Technologies, TDT) and demodulated into two signals using TDT’s Synapse software, corresponding to the 465 and 405 nm excitation wavelengths. To analyze the calcium activity, we used custom-written Python scripts. First, both signals were low-pass filtered at 2 Hz using a 4th order digital Butterworth filter. Next, using linear regression, we fitted the 405 nm to the 465 nm signal. Finally, the linear fit was subtracted from the 465 nm signal (to correct for photo-bleaching or motion artifacts) and the difference was divided by the linear fit yielding the ΔF/F signal. To determine the sleep-wake state, EEG and EMG signals were simultaneously recorded with calcium signals using the RZ5P amplifier.

To detect calcium transients occuring on the infraslow timescale, we first filtered the calcium signal with a zero-lag, 4th order digital Butterworth filter with cutoff frequency 1/15 Hz as described in^20^. Next, we detected prominent peaks in the signal using the function find_peaks provided by the python library scipy (https://scipy.org/). As parameter for the peak prominence, we used 0.05 * distance between the 1st and 99th percentile of the distribution of the ΔF/F signal.

### Analysis of infraslow σ power oscillations

To calculate the power spectral density of the EEG σ power, we first calculated for each recording the EEG power spectrogram by computing the FFT for consecutive sliding, half-overlapping 5 s windows. Next, we normalized the spectrogram by dividing each frequency component by its mean power and calculated the normalized σ power by averaging across the spectral density values in the σ range (10.5 - 16 Hz). As the infraslow rhythm is most pronounced in consolidated NREMs bouts^30^, we only considered NREMs bouts that lasted at least 120 s, possibly interrupted by MAs (wake periods ≦ 20s). We then calculated the power spectral density using Welch’s method with Hanning window for each consolidated NREMs bout and averaged for each animal across the resulting densities. To quantify the strength of the infraslow rhythm, we computed the areas under the PSD in the ranges 0.01 - 0.04 Hz and 0.08 - 0.12 Hz, respectively, and subtracted the second value from the first value.

### Sleep spindle detection

Spindles were detected using a previously described algorithm using the frontal EEG^28^. The spectrogram was computed for consecutive 600 ms windows with 500 ms overlap, resulting in a 100 ms temporal resolution. The spindle detection algorithm used two criteria to determine for each 100 ms time bin whether it was part of a spindle or not: The first criterion was that the height of the maximum peak in the σ frequency range (10 - 16.67 Hz) exceeds a threshold, which corresponded to the 96th percentile of all maximum peaks in the σ frequency range of the sleep recording. We determined the optimal percentile value by maximizing the performance of the algorithm on a manually annotated control data set. Second, the power value of the peak in the σ range (10 - 16.67 Hz) had to be greater than half of the peak value in the range 0 - 10 Hz. The optimal value for this ratio (σ peak ratio) was again determined on the control data set. Next, the algorithm merged spindle events that were temporally close to each other. First, spindle events in adjacent bins were considered as part of the same spindle. Second, we fused together sequences of spindle events that were interrupted by gaps of less than 300 ms. The optimal value for the gap was again determined on the control data set. Finally, we discarded spindles with duration ≦ 200 ms. Of all the potential spindles, we only considered those as spindles where for at least half of the time bins the peak frequency lied in the range of 10 - 16.7 Hz. The parameters of the spindle detection algorithm (σ percentile threshold, σ peak ratio, and minimum fusing distance) were optimized using a manually annotated data set.

### Patch clamp recordings

Mice were deeply anesthetized with ketamine/xylazine (200/20 mg/kg body weight) and decapitated. Brains were harvested and placed immediately in ice-cold cutting solution (92 mM *N*-methyl-D-glucamine, 2.5 mM KCl, 1.2 mM NaH_2_PO_4_, 30 mM NaHCO_3_, 20 mM HEPES, 25 mM glucose, 5 mM sodium L-ascorbate, 2 mM thiourea, 3 mM sodium pyruvate, 10 mM MgSO_4_ and 0.5 mM CaCl_2_) and continuously bubbled with 95% O_2_ and 5% CO_2_. 200 μm thick coronal sections were cut with a vibratome (Leica VT 1200S) and placed in artificial cerebrospinal fluid (ACSF: 126 mM NaCl, 2.5 mM KCl, 1.2 mM MgSO4, 2.4 mM CaCl2, 25 mM NaHCO3, 1.4 mM NaH2PO4, 11 mM glucose and 0.6 mM sodium L-ascorbate) and continuously bubbled with 95% O_2_ and 5% CO_2_. Slices were incubated at 31°C for 30 minutes and then at room temperature for 30 minutes. Brain slices were transferred into a recording chamber and perfused with oxygenated ACSF. TdTomato+ cells in the LC were located under a 40X water-immersion objective (Olympus BX61WI). Recording pipettes were pulled from borosilicate glass (Flaming-Brown puller, Sutter Instruments, P-97, tip resistance of 5–10 MΩ) and filled with pipette solution consisting of 120 mM potassium gluconate, 10 mM NaCl, 1 mM CaCl_2_, 10 mM EGTA, 10 mM HEPES, 5 mM Mg-ATP, 0.5 mM Na-GTP and 10 mM phosphocreatine. Whole-cell patch clamp recordings were controlled via an EPC-10 amplifier and Pulse v8.74 (HEKA Electronik). Firing patterns upon current injection were recorded under current clamp mode.

### Spatial object recognition task

During the habituation session (ZT 3-4, days 1 and 2), mice were habituated to the training context (13”x13” open field arena) for two five-minute sessions. A visual cue (a rectangle containing alternating black and white stripes) is attached to one wall of the open field to help the mice orient themselves within the open field. During the training session (ZT 3-5, day 3), mice were placed in the arena with two identical objects (two small glass bottles) for three five-minute training sessions. Immediately following the training session, mice were returned to their home cage for 5 hrs. In Fig. 5C, optogenetic manipulation was performed for 5 hrs in their home cage (ZT 3.5-10: the first mouse received optogenetic stimulation between ZT 3.5-8.5, and the last mouse received stimulation between ZT 5-10) before the test session. During the test session (ZT 8.5-10.5, day 3), mice were placed in the arena with the two familiar objects, one displaced to a new location, for one 10 minute test session.

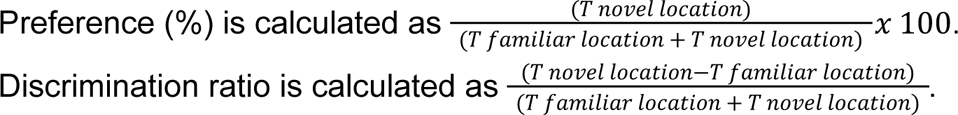

Mice that explored less than 2 s during test or at least two training sessions were excluded.

### Statistical tests

Statistical analyses were performed using the python packages scipy.stats (scipy.org) and pingouin (https://pingouin-stats.org)^54^. We did not predetermine sample sizes, but cohorts were similarly sized as in other relevant sleep studies^19,41^. All statistical tests were two-sided. Data were compared using t tests, bootstrap or ANOVA followed by multiple comparisons tests. For unpaired two sample T tests, Welch T test was used when the sample sizes are unequal as recommended^55^. For RM and mixed ANOVA, Mauchly’s test was applied to check the sphericity of the data. In case sphericity was violated, P values were corrected using the Greenhouse-Geisser correction. To account for multiple comparisons, P values were Bonferroni or Holm corrected. For all tests, a (corrected) P value < 0.05 was considered significant. Box plots were used to illustrate the distribution of data points. The upper and lower edges of the box correspond to the quartiles (25th and 75th percentile) of the dataset and the horizontal line in the box depicts the median, while the whiskers indicate the remaining distribution, except for outliers, i.e. points smaller than the 25th percentile - 1.5 * the interquartile range (IQR) or larger than the 75th percentile + 1.5 IQR. Outliers are depicted as diamonds. Statistical results and parameters (exact value of n and what n represents) are presented in the Table S1, figure legends and results.

## ACKNOWLEDGMENTS

This work was supported by the National Institutes of Health/National Institute of Neurological Disorders and Stroke (R01-NS-110865), Simons Foundation Pilot Award, Eagle Autism Challenge Pilot Grant, The Hartwell Individual Biomedical Research Award to S. C. We thank Mandy Schott for help with setting up sound stimulation experiments, and the members from Chung and Weber labs for helpful discussion.

## AUTHOR CONTRIBUTIONS

Conceptualization, A.C., and S.C.; Methodology, A.C., J.S., A.W., and J.P.B.; Software, H.A., F.W.; Investigation, A.C., J.S., Y.W., H.S., B.K., X.J., J.H., and I.A.; Resources, S.T., and K.B.; Writing, Review & Editing, A.C., M.M., F.W and S.C.; Visualization, A.C.; Supervision, S.C.; Funding Acquisition, S.C.

## DATA AND CODE AVAILABILITY

All datasets will be shared by the lead contact upon request after publication. All original code is deposited in: https://github.com/tortugar/Lab

